# Discovery of allosteric non-covalent KRAS inhibitors that bind with sub-micromolar affinity and disrupt effector binding

**DOI:** 10.1101/440487

**Authors:** Michael J. McCarthy, Cynthia V. Pagba, Priyanka Prakash, Ali Naji, Dharini van der Hoeven, Hong Liang, Amit K. Gupta, Yong Zhou, Kwang-Jin Cho, John F. Hancock, Alemayehu A. Gorfe

**Affiliations:** Department of Integrative Biology and Pharmacology, McGovern Medical School, The University of Texas Health Science Center at Houston, 6431 Fannin St., Texas 77030, USA; Biochemistry and Cell Biology Program, UTHealth MD Anderson Cancer Center Graduate School of Biomedical Sciences, Houston, 6431 Fannin St., Texas 77030, USA; Department of Diagnostic and Biomedical Sciences, School of Dentistry, The University of Texas Health Science Center at Houston, 7500 Cambridge St., Texas, USA; Department of Biochemistry and Molecular Biology, Boonshoft School of Medicine, Wright State University, Dayton, Ohio 45435, USA

## Abstract

Approximately 15% of all human tumors harbor mutant KRAS, a membrane-associated small GTPase and a notorious oncogene. Somatic mutations that render KRAS constitutively active lead to uncontrolled cell growth, survival, proliferation, and eventually cancer. KRAS is thus a critical anticancer drug target. However, despite aggressive efforts in recent years, there is no drug on the market that directly targets KRAS. In the current work, we combined molecular simulation and high-throughput virtual screening with a battery of cell-based and biophysical assays to discover a novel, pyrazolopyrimidine-based allosteric KRAS inhibitor that exhibits promising biochemical properties. The compound selectively binds to active KRAS with sub-micromolar affinity, slightly modulates exchange factor activity, disrupts effector Raf binding, significantly reduces signal transduction through mutant KRAS and inhibits cancer cell growth. Moreover, by studying two of its analogues, we identified key chemical features of the compound that are critical for affinity, effect on effector binding and mode of action. We propose a set of specific interactions with key residues at the switch regions of KRAS as critical for abrogating effector binding and reducing the rate of nucleotide exchange. Together, these findings not only demonstrate the viability of direct KRAS inhibition and offer guidance for future optimization efforts, but also show that pyrazolopyrimidine-based compounds may represent a first-in-class lead toward a clinically relevant targeting of KRAS by allosteric non-covalent inhibitors.

## Introduction

Somatic mutations in RAS proteins are associated with about 16% of all human cancers (1, 2). KRAS is the most frequently mutated RAS isoform, accounting for 85% of all RAS-related cancers (1, 2). Cellular KRAS is tethered to the inner surface of the plasma membrane by a farnesylated polybasic lipid anchor (3), and cycles between active guanosine tri-phosphate (GTP)- and inactive guanosine di-phosphate (GDP)-bound conformational states (4). GTPase activating proteins (GAPs) facilitate hydrolysis of GTP by KRAS while guanine nucleotide exchange factors (GEFs) catalyze GDP dissociation (4–6). Upon activation by receptor tyrosine kinases such as epidermal growth factor (EGF) receptors, GEFs are recruited to KRAS and initiate exchange of GDP for GTP. Active KRAS interacts with effectors such as Raf in the MAPK pathway and PI3K in the AKT pathway (7), driving cell growth and proliferation (8, 9). In a regulated RAS cycle, signaling is turned off upon GTP hydrolysis. Oncogenic mutations that impair its GAP-mediated or intrinsic GTPase activity render KRAS constitutively active and thereby cause uncontrolled cell growth/proliferation leading to cancer (1, 2). Mutant KRAS is therefore a highly sought-after anticancer drug target (10, 11).

Despite decades of efforts, however, drugging KRAS (and RAS proteins in general) remains an unrealized goal (12). Among the many challenges, conservation of the nucleotide-binding site among a diverse group of small GTPases (4, 13), and the high (picomolar) affinity of RAS for its endogenous ligands GDP or GTP, are arguably the most significant. These issues made competitive inhibition impractical and avoiding off-target effects difficult. Thus, along with efforts at indirect RAS inhibition by targeting its interaction partner proteins (14, 15) or membrane localization (16, 17), development of direct allosteric KRAS inhibitors is currently a major focus of many laboratories (18). Proof-of-principle studies have established the allosteric nature of RAS (11, 19, 20), and discovered several allosteric small-molecule KRAS binders (21–25). Moreover, a number of recent reports described molecular fragments (23), small-molecules (18, 24–26), peptidomimetics (27, 28) and monobodies (29) that bind KRAS and modulate its functions in various ways. While this paints an optimistic picture of the prospects of allosteric KRAS inhibition, to the best of our knowledge none of these compounds has made it to clinical trial. Recent efforts toward developing covalent GDP analogues (30) or other small-molecule ligands (31) targeting G12C mutant KRAS may have a better chance of eventually treating specific tumor types (18). However, their application is likely limited to a few cancer cases such as small cell lung cancer (10). We believe non-covalent allosteric inhibition will be needed to target some of the most important mutations in KRAS including G12D, G12V, G13D and Q61H that are critical in biliary tract, small intestine, colorectal, lung and pancreas cancers (2, 10). Together, these four mutations appear to account for greater than 78% of all KRAS-associated cancers (10).

In previous reports, we described four allosteric ligand-binding sites on KRAS using a range of computational approaches (32, 33), including molecular dynamics (MD) simulations to sample transient conformations with open allosteric pockets (34–36). Among these, pocket p1 was the best characterized and is well established as a suitable target with many crystal structures of p1-bound ligand-KRAS complexes available in the protein data bank (PDB). In the current work, we combined MD simulation with a range of biophysical and cell assays to discover and characterize a novel class of inhibitors that bind to the p1 pocket with sub-micromolar affinity and abrogate signaling primarily by directly inhibiting the interaction of KRAS with effector proteins.

## Materials and Methods

### Molecular dynamics simulation and allosteric pocket analysis

Most oncogenic RAS mutants are constitutively active because their ability to hydrolyze GTP is compromised (37, 38). An inhibitor that selectively targets GTP-bound mutant RAS would therefore be desirable. However, there was no ligand-free high-resolution experimental structure of GTP-bound KRAS (^GTP^KRAS) when we started this project in 2014, and our target pocket p1 (see below) was closed or was too small in the available GDP-bound KRAS (^GDP^KRAS) structures. Therefore, we used MD simulation to generate an ensemble of ^GTP^KRAS structures with open p1. The initial structure for the simulation was a 5’-guanosinediphosohate-monothiophosphate (GSP)-bound KRAS^G12D^ X-ray structure from the PDB (ID 4DSO) with benzamidine bound at p1 and glycerol between helices 2 and 3 (23). After converting GSP to GTP, removing all other molecules except crystal waters and the bound Mg^2+^, adding hydrogen atoms and solvent, minimization and restrained simulation, we conducted a 300 ns production run using an identical protocol to that described in a recent report (39). The trajectory was analyzed in terms of volume and other features (such as numbers of hydrogen bond donors and acceptors) of our target pocket p1, and the conformation with the most open p1 was selected for virtual screening of ligand libraries.

### High throughput virtual screening

Six million compounds from the Drugs Now subset of the ZINC (40) database were docked into pocket p1 of our MD-derived KRAS^G12D^ structure (Fig 1A, see also Fig S1). Gasteiger charges and atomic radii were assigned using AutoDock Tools, and a first round of docking was conducted with AutoDock (41) as implemented in the parallelization routine DOVIS (42). We used the flexible ligand option with 1.0 Å spacing, along with a Lamarckian search with 150 generations and 1,000,000 energy evaluations. The top ~4000 compounds with energy score ≤ −6.8 kcal/mol were re-screened with VINA v1.1.2 (43) with exhaustiveness set to 12 and energy range to 4. The top 500 hits in each screen were then evaluated in terms of their ability to form close contact, salt bridge, hydrogen bonding, hydrophobic, cation-π, π-π and π-stacking interactions with the protein, using distance and angle cutoffs recommended by Durant et. al. (44). We found 58 ligands that score well in the majority of these metrics and experimentally tested 11 that are listed in Fig S2A.

### Cell signaling

The inhibitory potential of compounds was tested in monoclonal Baby Hamster Kidney (BHK) cell lines stably expressing monomeric green fluorescence protein (mGFP)-tagged KRAS^G12D^, KRAS^G12V^ and HRAS^G12V^. Cells were cultured in Dulbecco’s Modified Eagle Medium (DMEM, Hyclone) supplemented with 10% v/v bovine calf serum and incubated with compound or vehicle (DMSO) for 3 h without serum. Cells were then harvested in lysis buffer (50 mM Tris (pH 7.5), 75 mM NaCl, 25 mM NaF, 5 mM MgCl_2_, 5 mM EGTA, 1 mM dithiothreitol, 100 μM NaVO_4_, 1% Nonidet P40 plus protease inhibitors) and subjected to Western analysis controlling protein loading by BCA (bicinchoninic acid) assay. Lysates were resolved with BioRad polyacrylamide TGX 10% gel, transferred to polyvinylidene fluoride (PVDF) membrane and immunoblotted using pan-AKT (2920S), GFP (2956S), p-AKT^S473^ (4060L), p-cRaf^S338^ (9427S), p-ERK^T202/Y204^ (4370L), ERK1/2 (4695S) or β-actin antibodies (Cell Signaling Technology). IC_50_ values were calculated with Prism 4-parameter fit.

### Pull-down

We pulled down GFP-RAS with GST-tagged RAS binding domain (RBD) of cRaf^A85K^ (hereafter GST-Raf^RBD^) to monitor RAS-Raf interaction. To prepare GST-Raf^RBD^ bound to agarose beads, bacteria (BL21) transfected with a previously cloned GEX plasmid were grown in selection media to OD levels of 0.5-1.0 before protein expression was initiated with IPTG (1:1000). After 4 h, the sample was centrifuged at 6000 rpm for 5 min at 4°C, the pellet was resuspended with PBS containing 5 mM EGTA, 1% Triton X-100, PIC 1:50 and PMSF 1:100, and cells were lysed with cycles of freezing and thawing. The lysate was sonicated to breakup DNA, and pelleted. The supernatant was incubated with glutathione agarose (Pierce™) beads that bind to GST-Raf^RBD^. For all pull-down experiments, equal volumes of lysates from BHK cells expressing GFP-RAS were incubated for 2 h at 4°C with GST-RBD beads plus control DMSO or compound. Then samples were washed with Tris buffer (50 mM Tris, pH 7.4, 1 mM EDTA, 1 mM EGTA, 150 mM NaCl, 0.1% Trition X-100 and protease inhibitors) and immunoblotted with anti-GFP (Cell Signaling) and anti-GST (Santa Cruz Biotechnology) antibodies.

### Fluorescence lifetime imaging (FLIM)-fluorescence resonance energy transfer (FRET)

FLIM-FRET experiments were carried out using a lifetime fluorescence imaging attachment (Lambert Instruments, The Netherlands) on an inverted microscope (45). BHK cells transiently expressing mGFP-tagged KRAS^G12D^ (donor), alone or with mRFP-tagged cRaf^WT^ (acceptor) (1:5 ratio), were prepared and treated with compound for 2 h, washed with PBS, fixed with 4% paraformaldehyde (PFA) and quenched with 50 mM NH_4_Cl. The samples were excited using a sinusoidally modulated 3 W 470 nm light-emitting diode at 40 MHz under epi-illumination. Fluorescein (lifetime = 4 ns) was used as a lifetime reference standard. Cells were imaged with a Plan Apo 60X 1.40 oil objective using an appropriate GFP filter set. The phase and modulation were determined from 12 phase settings using the manufacturer’s software. Resolution of two lifetimes in the frequency domain was performed using a graphical method (46) mathematically identical to global analysis algorithms (47, 48). The analysis yields the mGFP lifetime of free mGFP donor (τ1), and the mGFP lifetime in donor/acceptor complexes (τ2). FLIM data were averaged on a per-cell basis. In a separate set of experiments, BHK cells co-expressing GFP-KRAS^G12D^ or GFP-HRAS^G12V^ with empty vector pC1 or mCherry-RBD were treated with vehicle DMSO or 1 uM and GFP fluorescence was measured as described above.

### Cell proliferation

Potential effect of the ligands on cancer cell proliferation was tested in four lung cancer cells: SKLU-1 (KRAS^WT^), H1975 (KRAS^WT^), H441 (KRAS^G12V^) and H522 (KRAS^G12D^), and four oral cancer cell lines: UM-SCC-22A (HRAS^WT^), UM-SCC-22A (HRAS^G12V^), HN31 (HRAS^G12D^) and HN31 (HRAS^knockdown^). 1000 cells were seeded per well in a 96-well plate. After 24 h of seeding, fresh growth medium supplemented with vehicle (DMSO) or varying concentrations of drug was added. Cells were treated with drug for 72 h, with addition of fresh medium containing drug every 24 h. Then cells were washed with PBS and frozen at −80°C for a minimum of 24 h. Plates were thawed and CyQuant™ dye (in lysis buffer provided in the CyQuant™ cell proliferation assay kit, Invitrogen™) was added. After a five-minute incubation, fluorescence (excitation: 480 nm emission: 520 nm) was measured with a Tecan Infinite M200 plate reader for the lung cancer cells, and numbers of the oral cancer cells were quantified using the CyQuant Poliferation Assay (ThermoFisher), according to manufacturer’s protocol.

### Microscale thermophoresis (MST)

Determination of dissociation constants using MST was performed following vendor protocols. Purified KRAS was labeled with the Monolith MT™ Protein Labeling Kit RED-NHS (NanoTemper Tech) through buffer-exchange in the labeling buffer (40 mM HEPES, pH 7.5, 5 mM MgCl_2_, 500 mM NaCl). The concentration of the eluted protein was adjusted to 2-20 μM, the dye added at a 2-3-fold concentration to a final volume of 200 μL, and the mixture incubated for 30 min at room temperature in the dark. Labeled KRAS was purified using the column provided in the kit. For MST measurements, a 16-point serial dilution of ligand was prepared in an MST assay buffer (40 mM HEPES, pH 7.5, 5 mM MgCl_2_, 100 mM NaCl, plus 0.05% Tween-20 and 2-4% DMSO), and added to an equal volume of a 100 nM KRAS solution. The solutions were loaded in capillaries and measurements were done at room temperature using 20% LED and 40% MST power. The data were fit in Igor Pro using the Hill equation.

### Nucleotide exchange and release assays

Loading of fluorescent-labeled GDP (BODIPY-GDP; BGDP from hereon) to KRAS was conducted following previous reports (23, 49), with minor modifications. Purified KRAS was buffer-exchanged in NAP-5 column (GE Life Sciences) in low Mg^2+^ buffer (25 mM Tris pH 7.5, 50 mM NaCl, and 0.5 mM MgCl_2_). The eluate was incubated with 10-fold molar excess of BGDP (Life Technologies) in the presence of 5 mM EDTA and 1 mM DTT for 1.5 h at 20 °C in the dark. Then 10 mM MgCl_2_ was added and the solution was incubated for 30 min at 20 °C. Free nucleotide was removed by gel filtration using a PD-10 column (GE Life Sciences) that had been equilibrated with the reaction buffer (25 mM Tris-HCL pH 7.5, 50 mM NaCl, 1 mM MgCl_2_, and 1 mM DTT). The concentration of ^BGDP^KRAS was determined using Bradford assay and a BGDP standard curve. Then the effect of ligands on the intrinsic rate of nucleotide release was monitored using the decrease in fluorescence with time as BGDP dissociates from KRAS in a 100 μL reaction mixture (96-well plate) of 0.5 μM ^BGDP^KRAS, 100 μM GTP and varying concentrations of ligand (0 to 25 μM); GTP was added just before measurements. To measure the rate of SOS-mediated nucleotide release, 0.5 μM SOS (residues 564-1049, Cytoskeleton Inc) was added after adding GTP, and fluorescence was immediately read (excitation: 485 nm, emission: 510 nm) using Tecan Infinite M200 plate reader. Intrinsic and SOS-mediated nucleotide exchange rates were monitored with the fluorescence intensity increase of BGTP as it displaces GDP from KRAS. We used a 100 μL reaction mixture containing 0.5 μM each of ^GDP^KRAS, BGTP (and SOS) plus varying concentrations of ligand (0 to 25 μM); BGTP was added just before measurements. Experiments were conducted with minimal light and the reaction was monitored for 2 h at room temperature. Fluorescence intensities were normalized at 120 s and the traces were fit with linear or single exponential functions (Igor Pro, Wavemetrics).

### Fluorescence polarization

Fluorescence polarization assay was conducted following previous reports (50, 51). KRAS was pre-loaded with the non-hydrolyzable fluorescent GTP analog BODIPY-GTP-γ-S (BGTP-γ-S; Life Technologies) using buffer-exchange in NAP-5 (GE Life Sciences) as described in the previous section. Then 0.5 μM (50 μL) of ^BGTP-γ-S^KRAS was incubated with an equal volume but varying concentrations (0 to 2.5 μM) of GST-Raf^RBD^ (Raf RBD residues 1-149; Life Technologies) for 30 min in the dark. To determine the effect of ligand on RAS-Raf binding, KRAS was first incubated with a fixed concentration of the ligand for 30 min and then with GST-Raf^RBD^. Fluorescence polarization was measured using PolarStar Optima plate reader (excitation: 485 nm, emission: 520 nm) at room temperature. GST-tag was used to increase the weight of Raf^RBD^ for a greater polarization. The dissociation constant for KRAS-Raf binding was determined using a quadratic ligand binding equation (50).

## Results and Discussion

### Initial hits from molecular modeling and high-throughput virtual screening

We conducted *in silico* screening of compounds from the ZINC database (40) targeting pocket p1 on an MD-derived structure of ^GTP^KRAS^G12D^. This pocket is located between the functionally critical switches 1 (residues 25-40) and 2 (residues 60-75), and encompasses residues 5-7, 37-39, 50-56, 67 and 70-75 (Figs 1A, S1). Many of these residues, including residues 37-39 on the effector binding loop and residue 71 on switch 2, participate in interaction with effectors and/or GEFs. Therefore, we reasoned that a p1-targeted ligand could disrupt either or both of these interactions. However, p1 was fully or partially closed in the available KRAS structures including the holo forms, which were generally bound to small (<160 Da) ligands (Fig S1). We wanted to have a more open conformation in order to dock a wide range of “drug-like” molecules spanning the ~150-500 Da molecular weight common in marketed drugs. We therefore conducted MD simulation to generate an ensemble of ^GTP^KRAS^G12D^ structures with open p1. Analysis of the trajectory yielded 119 and 219 Å^3^ as the mean and maximum volumes of pocket p1, respectively. We performed retrospective comparison of the MD conformer with the most open p1, which we used for molecular docking, with currently available GTP (or analogue)- and GDP-bound crystallographic KRAS structures (Fig S1). We observed three distinct groups of conformers that differ mainly in the orientation of helix 2. In one group, the orientation of helix 2 is such that pocket p1 is nearly or completely closed (orange). All of these structures are GDP-bound and are dominated by structures in complex with covalent ligands. In the second, sampled by both GDP-and GTP-bound KRAS, movement of helix 2 toward helix 3 opens up the pocket to some extent. In group 3, helix 2 moved even farther away from the core beta-sheet, allowing for a more open p1. Our MD-derived conformer belongs to the third group and exhibits the largest displacement of helix 2, which, together with side chain re-orientations, allowed for a wider pocket p1 (Fig S1). We used this snapshot to conduct an initial screen of 6,000,000 compounds, followed by a secondary screen of the top ~4000 (see Methods). Analysis of the top 500 ligands in each screen yielded a consensus prediction of 58 initial hits. Eleven of these were purchased and tested in cells (Fig S2A).

**Figure 1.**
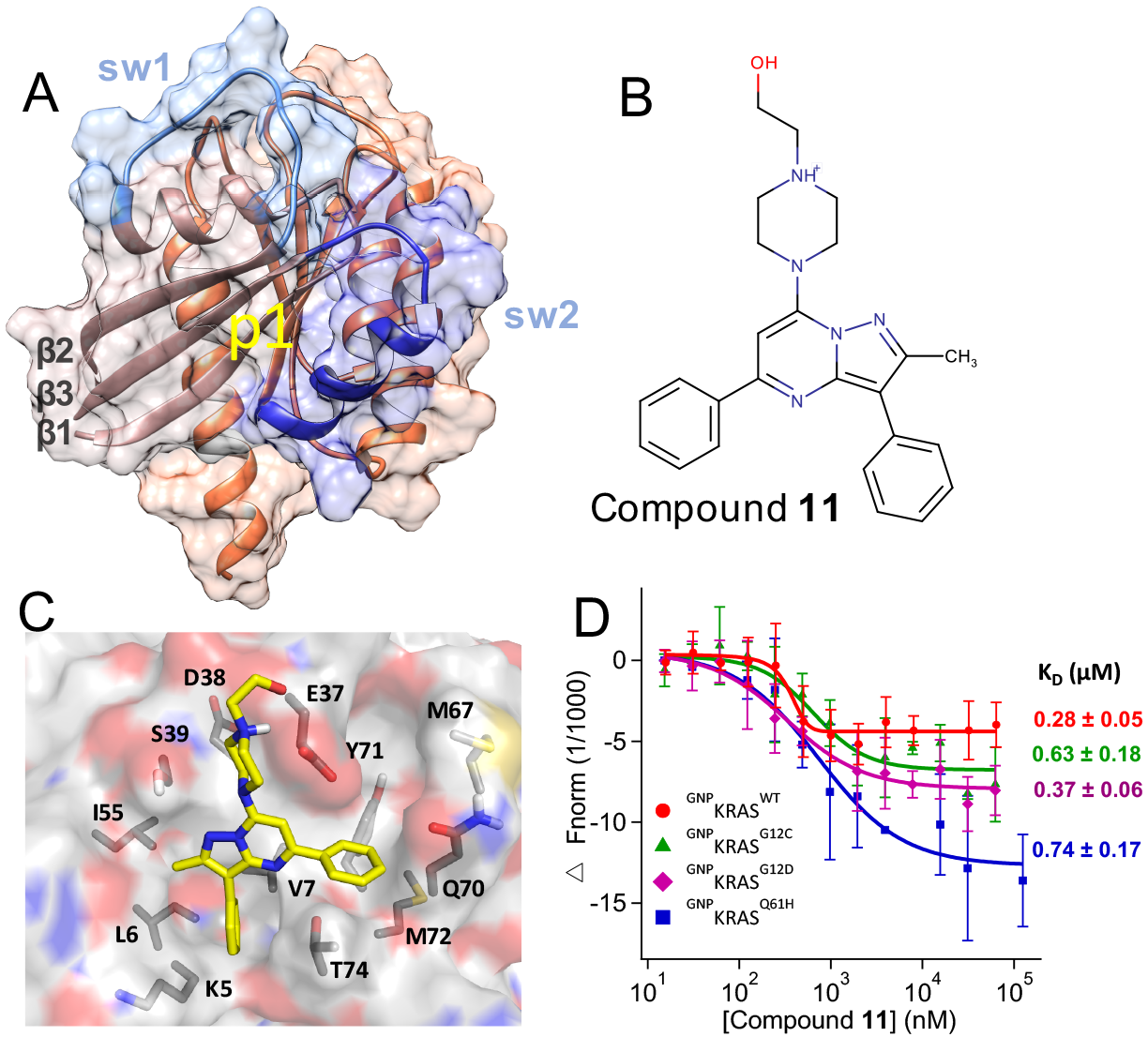
Predicted binding mode and measured affinity of compound 11 to KRAS. (**A**) Structure of the catalytic domain of KRAS used for the virtual screening. Lobe1 (residues 1-86) and lobe2 (residues 87-166) are highlighted in different colors, as are switch 1 (residues 30-40) and 2 (residues 60-75). The location of our target allosteric pocket p1 is indicated. (**B**) Chemical structure of compound **11**. **(C)** Predicted binding pose of compound **11**, with the key residues that make polar or vdW contacts with the ligand labeled. **(D)** Microscale thermophoresis (MST) experiments indicating direct binding of compound **11** to KRAS, along with dissociation constants (K_D_) derived from the curves. Changes in fluorescence upon titration of 50 nM KRAS with increasing concentration of compound are shown: KRAS^WT^ (red), KRAS^G12C^ (green), KRAS^G12D^ (purple) and KRAS^Q61H^ (blue), each bound to the non-hydrolyzable GTP analogue, guanylyl imidodiphosphate (GNP). No or very weak binding was detected to GDP-bound KRAS, GNP- or GDP-bound NRAS and HRAS, and Rap1B which was used as control (Fig S3).

### Cell signaling assays identify compound 11 as a promising initial hit

Western analysis was used to quickly assess the potential impact of our predicted hits on MAPK signaling, a major pathway mediated by KRAS. Specifically, we monitored ERK1/2 phosphorylation levels (p-ERK) in BHK cells stably expressing KRAS^G12D^ treated with vehicle (DMSO), the MEK inhibitor U0125 (U) or compound at four different concentrations (1-100 μM). The results showed that the majority of the predicted hits have no effect while few (e.g. **4**) increase rather than decrease p-ERK levels (Fig S2B). Compounds **9** and **11**, on the other hand, decreased p-ERK levels at concentrations ≥ 50 μM and ≥1 μM, respectively. To verify the latter observation, we repeated the experiments in an expanded concentration range starting from 0.1 μM. As in the first screen, compound **11** dose-dependently decreased p-ERK levels, leading to a ~50% reduction at 5 μM (Fig S2C). However, compound **9** increased p-ERK levels at 25 and 38 μM in contrast to the decrease observed at higher concentrations (Fig S2B). Although a similar increase and then decrease of KRAS signaling upon increasing ligand concentration has been observed before (49, 52), we selected the more potent and monotonously dose-dependent compound **11** for further analysis.

### Compound 11 binds to WT and oncogenic KRAS mutants with high affinity

Figures 1B and 1C show the chemical structure and the predicted complex of compound **11** with KRAS, suggesting that the ligand potentially forms multiple favorable interactions with residues in the p1 pocket. Figure 1D shows that the compound binds to the catalytic domain (residues 1-166) ^GTP^KRAS^WT^ with a K_D_ = ~ 0.3 μM, suggesting a very tight binding rarely seen in primary screens. The compound has a very similar affinity (K_D_ = ~0.4-0.7 μM) for oncogenic mutants KRAS^G12D^, KRAS^G12C^ and KRAS^Q61H^ in the GTP state (Fig 1D). However, no binding or very weak binding was detected for KRAS^WT^ and KRAS^G12D^ in the GDP state, HRAS^WT^ and NRAS^WT^ in both their GDP and GTP-bound forms, or to our control Rap1b (Fig S3), a RAS-related small GTPase with homologous structure. Few weak-affinity non-covalent binders that exhibit selectivity toward GDP- or GTP KRAS have been reported (23–25). However, to the best of our knowledge, compound **11** is the first small molecule to selectively bind to KRAS^GTP^ with a nanomolar affinity.

In the docked pose (Fig 1C), the 1-piperazineethanol moiety occupies an electronegative cleft near D54 and D38, potentially donating hydrogen bonds to the side chain and backbone atoms of E37. The methylated pyrazolopyrimidine core sits in a trench on top of V7 and L56 with its methyl group pointing towards I55 while the pyrimidine-bound benzene ring occupies the space between the central beta sheet (β1-β3) and helix 2, and makes π-stacking interaction with Y71. The pyrazol-attached benzene is buried deep in a tight pocket, stabilized primarily by van der Waals (vdW) interactions with side chain carbon atoms of V7, L6 and K5. These interactions are common in the majority of our predicted hits listed in Fig S2A. Therefore, we propose that, in addition to potential induced-fit effects, compound **11**’s preference for GTP-bound KRAS may be due to conformational differences of these residues in ^GTP^RAS versus ^GDP^RAS (4). Comparison of available GDP- and GTP-bound RAS structures supports this conclusion. For example, pocket p1 is partially occluded by helix 2 in a large number of GDP-bound KRAS (Fig S1) and HRAS (Fig S4) crystallographic structures. Similarly, compound **11**’s apparent preference for KRAS over HRAS or NRAS may arise from subtle differences in structure or dynamics. For example, Mattos and colleagues have recently shown that the active site of activated KRAS is more open and dynamic than that of HRAS (53).

### Compound 11 disrupts interaction of KRAS with Raf

We used three different assays to check if our compound inhibits RAS signaling by interfering with effector binding. These included fluorescence polarization and pull-down assays, which directly measure the interaction of KRAS with the RBD of Raf in purified or cell lysate systems, respectively, and FLIM-FRET, which measures the interaction of KRAS with full-length or truncated Raf in the cellular milieu. We used fluorescence polarization of BGTP-γ-S to monitor binding of the KRAS catalytic domain to GST-Raf^RBD^ with and without pre-incubation with 1 μM compound **11**. Fig 2A shows a dramatic decrease in polarization in the entire concentration range of GST-Raf^RBD^. For example, at 2 μM GST-Raf^RBD^, compound treatment reduced the polarization and therefore RAS-Raf^RBD^ interaction by >80%. That we observed such a large reduction despite the weaker affinity of the RBD used in this assay (residues 1-149) than the commonly used shorter RBD (residues 51-131) further highlights the major impact of **11** on KRAS/Raf complex formation. The dissociation constant derived from the polarization curves indicate that **11** reduced the affinity of KRAS to Ra1^RBD^ by ~13-fold. Consistent with this observation, pull-down of GFP-KRAS^G12D^ by GST-Raf^RBD^ in compound-treated cell lysates show a significant (e.g. >50% at 1 μM of **11**) decrease in GFP-KRAS^G12D^ levels (Fig 2B).

We observed a similar effect in FLIM-FRET experiments in cells. In this experiment, quenching of GFP fluorescence lifetime indicates RAS-cRaf interaction in cells co-transfected with GFP-RAS and RFP-cRaf. In cells co-expressing KRAS^G12D^ and wild type full-length cRaf, quenching of GFP fluorescence lifetime and hence KRAS^G12D^-cRaf interaction is significantly reduced upon compound treatment (Fig 2C). FLIM-FRET was also used to examine interaction of GFP-tagged RAS mutants and mCherry-tagged Raf^RBD^. As shown in Fig 2D, GFP fluorescence lifetime in cells expressing GFP-KRAS^G12D^ with empty vector pC1 was ~2.3 ns, which decreased to ~1.93 ns in cells co-expressing GFP-KRAS^G12D^ and mCherry-RBD, indicating significant FRET and thus interaction between the two constructs. Treatment with 1 μM of **11** for 2 h increased the GFP lifetime to ~2.02 ns, suggesting reduction of the interaction between KRAS^G12D^ and RBD. The same experiments with GFP-HRAS^G12V^ and mCherry-RBD show that compound **11** has the opposite, albeit smaller, effect on the interaction of HRAS^G12V^ with Raf^RBD^. These results in cells confirm our observations from pull-downs in lysates and fluorescence polarization in purified systems, as well as the KRAS-selectivity of compound **11** suggested by MST.

**Figure 2.**
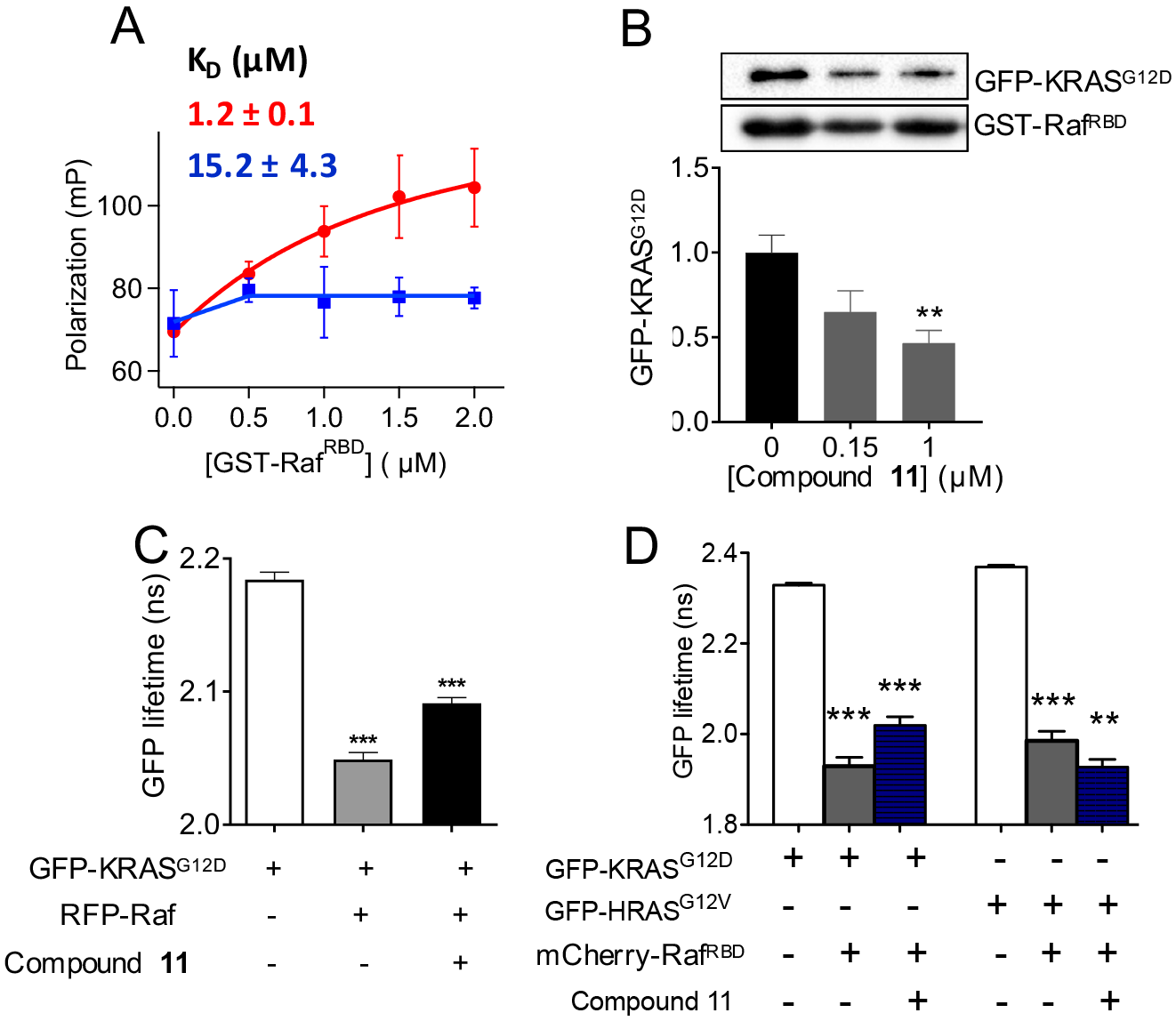
Compound 11 disrupts KRAS-Raf interaction. (**A**) Fluorescence polarization of ^BGTP-γ-S^KRAS (0.5 μM) as a function of a varying concentration of GST-Raf^RBD^ in the absence (red) and presence (blue) of 1 μM compound **11**. Shown above the curves is the K_D_ for KRAS-Raf^RBD^ binding obtained by fitting the data to 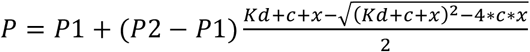, where *P1* is polarization of free KRAS, *P2* is polarization of Raf-bound KRAS, *c* is the total concentration of KRAS, and *x* the total concentration of Raf^RBD^. **(B)** Amount of GFP-KRAS^G12D^ pulled down by GST-Raf^RBD^ after treatment of cell lysates with compound at the concentrations indicated (representative Westerns shown at the top). An equal volume of lysates was used, and the data were normalized to GST-RBD and DMSO control which also serves as loading control. The RBD sequence length was 1-149 and 51-131 in the fluorescence polarization and pull-down assays, respectively. While the shorter RBD is sufficient for biochemical assays, the extra amino acids in the longer RBD increases the size and thereby enhance signal to noise ratio in the fluorescence polarization assay. **(C)** GFP fluorescence lifetime from FLIM-FRET using cells expressing GFP-KRAS^G12D^ alone or with RFP-Raf (full-length cRaf: residues 1-648), with or without treatment by 1 μM compound **11**. **(D)** GFP fluorescence lifetime from FLIM-FRET using cells expressing GFP-KRAS^G12D^ or HRAS^G12V^ with an empty vector pC1 or mCherry-Raf^RBD^ (RBD: residues 51-131), with or without treatment with 1 μM compound **11**. In **(B-D)** data are shown as mean ± S.E from three separate experiments; significance was estimated by oneway ANOVA relative to the control for each bar in **B**, the 2^nd^ bar in **C**, and the 2^nd^ and 4^th^ bars in D, or relative to the bar immediately to the left of bar 3 in **C** and bars 3 and 6 in **D**.

### Compound 11 inhibits KRAS signaling

Fig 3 shows that compound **11** dose-dependently decreases both p-ERK and p-cRaf levels in BHK cells expressing KRAS^G12D^ and KRAS^G12V^, suggesting inhibition of RAS signaling via the MAPK pathway. The data also indicate that the ligand has a slightly lower IC_50_ for its direct effector cRaf (e.g., 0.7 μM in the case of KRAS^G12D^) than the two-steps removed ERK (1.3 μM). Note also that the IC_50_ for cRaf is very close to the K_D_ of the ligand for ^GTP^KRAS. Changes in phosphorylated AKT (p-AKT) levels show that the compound also inhibits signaling through the AKT pathway but to a lesser extent than the MAPK pathway. Together, these results suggest that the ligand disrupts MAPK signaling by acting on RAS or its upstream modulators. To test if compound **11** is selective for the KRAS isoform, we measured p-ERK and p-cRaf levels in BHK cells expressing the constitutively active HRAS^G12V^ (Fig 3, right). We found no significant effect on the phosphorylation of these effectors and hence signaling via the MAPK pathway. Similarly, no statistically significant effect on p-AKT levels was observed even though H-Ras is a major driver of the AKT pathway. As a control, treatment of the HRAS^G12V^-expressing BHK cells with a 10 μM of our control U (the MEK inhibitor U0126) almost completely abolished MAPK signaling (Fig 3). We tested compound **11**’s effect on the proliferation of four lung and four oral cancer cell lines and found that the KRAS-expressing lung cancer cells, particularly those with mutant KRAS, are more sensitive to the compound than the HRAS-expressing oral cancer cells (Fig 4A). Also, there is no major difference between HRAS^WT^ and HRAS^G12V^/HRAS^G12D^ cancer cells, or between HN31 cancer cells with and without HRAS knockdown. To further illustrate that compound **11** inhibits the lung cancer cells with mutant KRAS more effectively, we plotted the relative growth of the eight cell lines in the presence of 5 μM compound **11** (Fig 4B). Relative to DMSO control, growth of the oral cancer cells with or without mutant HRAS, as well as the lung cancer cells with wild type KRAS, is 60-80% whereas the corresponding number for the lung cancer cells with KRAS^G12D^ and KRAS^G12V^ is 30-35%. In summary, the cell signaling and proliferation assays indicate that compound **11** selectively inhibits signaling through mutant KRAS, consistent with its significant effect on KRAS-Raf interaction (Fig 2).

**Figure 3.**
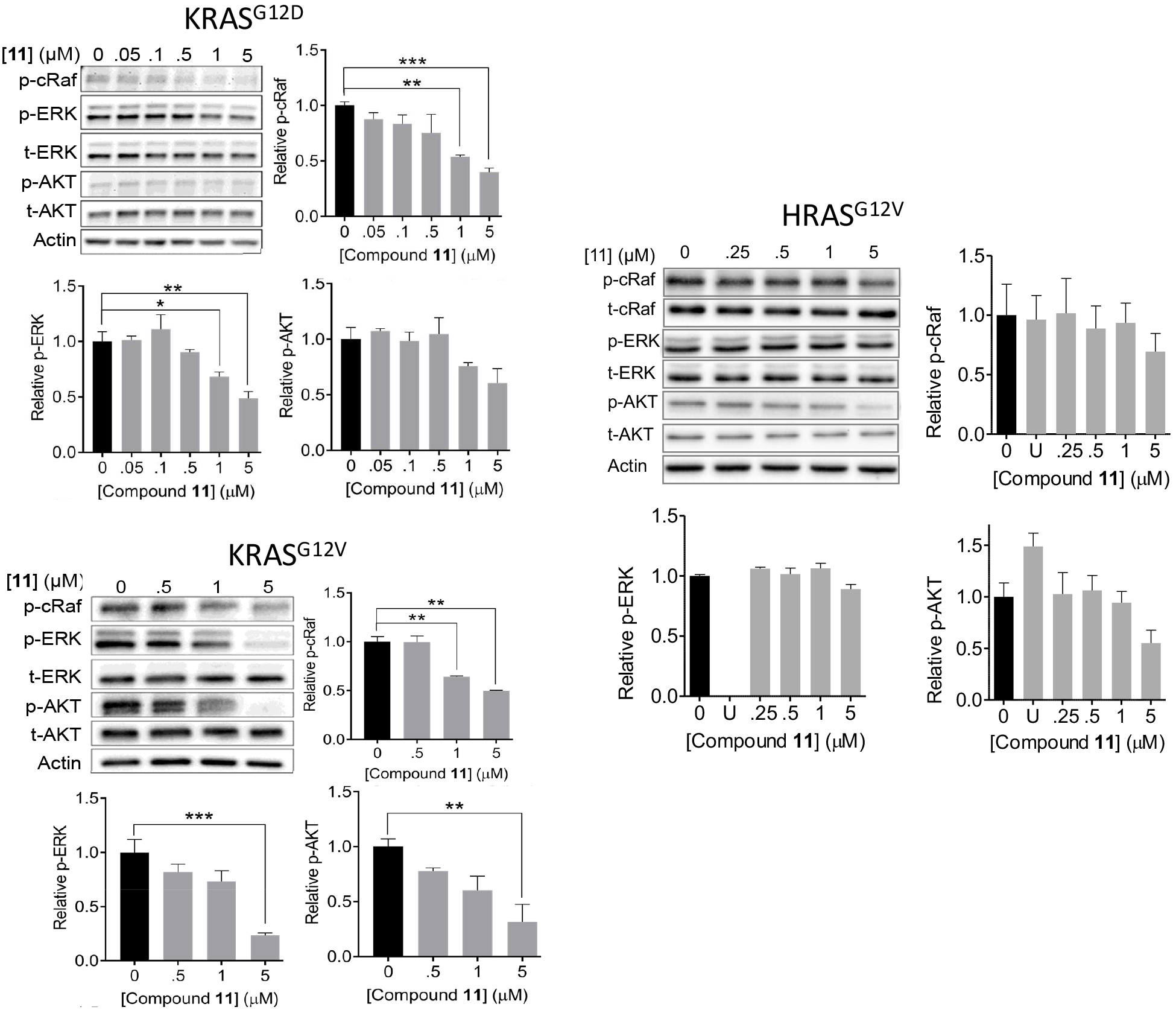
Compound 11 inhibits mutant KRAS signaling. Representative Western blots and their quantification showing levels of phosphorylated cRaf (p-cRaf), ERK (p-ERK) and AKT (p-AKT) in cells expressing KRAS^G12D^ (**top left**), KRAS^G12V^ (**bottom left**) and HRAS^G12V^ (**right**) treated with the indicated concentrations of compound **11**, DMSO or, where indicated 10 μM MEK inhibitor U0125 (U). Data are shown as mean ± S.E; significance was estimated by oneway analysis of variance: * = p < 0.02; ** = p < 0.005; *** = p < 0.0001.

**Figure 4.**
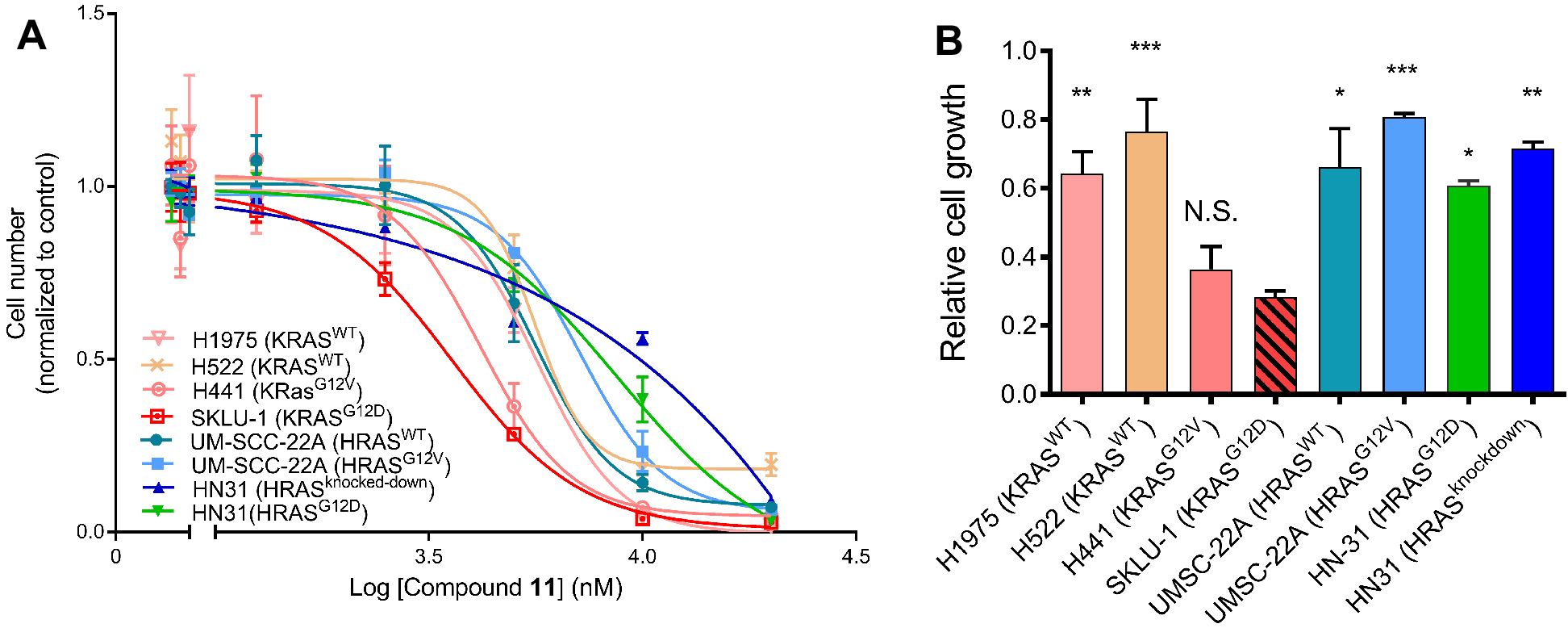
Cell proliferation assays suggest that cancer cells expressing mutant KRAS are more sensitive to compound 11. **(A)** Proliferation profile of KRAS-expressing lung cancer cells and HRAS-expressing oral cancer cells upon treatment by increasing concentration of compound **11** and monitored by CyQuant assay. **(B)** Relative growth of the KRAS and HRAS cancer cells after treatment with 5 μM of compound **11**. The lung cancer cells include H1975 and H522 that express KRAS^WT^, SKLU-1 that expresses KRAS^G12D^, and H441 harboring KRAS^G12V^. The oral cancer cells include UM-SCC-22A lines harboring HRAS^WT^ and HRAS^G12V^, HN31 cells expressing HRAS^G12D^, and HN31 cells with HRAS knockdown. Data are shown as mean ± S.E; significance was estimated by one-way analysis of variance with respect to the data for SKLU-1: * = p < 0.02; ** = p < 0.005; *** = p < 0.0001.

### Mechanism of action and optimization route for pyrazolopyrimidine-based KRAS inhibitors

In addition to its effect on effector binding, compound **11** also slightly reduced the rates of both intrinsic and SOS-mediated GDP/GTP exchange reactions of KRAS, as well as SOS-mediated GDP release (SI text and Fig S5). To identify the chemical fingerprints of compound **11** responsible for its high-affinity binding and effect on KRAS function, we studied compounds **12** and **13.** Obtained from similarity searches based on **11**, these analogues provided valuable insights into the mechanism of action of our pyrazolopyrimidine-based ligand. In compound **12**, the 1-piperazineethanol functional group of **11** is replaced by 1-methylpiperazine (Fig 5A,B), making it more hydrophobic and less soluble in DMSO. This compound slightly reduced p-ERK levels at a higher concentration of 2 μM (Fig 5C) but it is nearly as effective as **11** in inhibiting proliferation of lung cancer cells (Fig 5D). However, it has no effect on p-cRaf levels (Fig 5C) or on KRAS-Raf interaction as assessed by FLIM-FRET (Fig 5E), suggesting a potentially different mechanism of inhibition than compound **11**, or an off-target effect. The predicted binding mode of **12** is similar to that of **11**, but it lacks the capacity for hydrogen bonding interactions with residues at the effector-binding loop (Fig 5B). Together, these results suggest that the hydroxymethyl group on the piperazine ring, which in compound **11** is predicted to interact with residues in the effector-binding loop (Fig 1C), plays a crucial role in disrupting KRAS-Raf interaction and/or in modulating binding to KRAS.

Despite its several attractive features, solubility issues made compound **12** difficult to work with; a compound with a better solubility profile would be desirable. Also, we believe a derivative that preserves **11**’s effectiveness in inhibiting effector binding would make for a better lead compound. As shown below, compound **13** (Fig 6A,B) satisfies both of these conditions. It has a methyl group attached to the pyrimidine in place of the benzene ring found in **11**, which makes it less hydrophobic and readily soluble in DMSO and other common solvents. Therefore, we measured the K_D_ of the interaction of compound **13** with G12D and other KRAS mutants using MST. The results summarized in Fig 6C show that this compound has a 6.5-7.1-fold weaker affinity for KRAS than compound **11**. Similar to compound **11**, however, **13** does not bind to ^GDP^KRAS^WT^ or ^GDP^KRAS^G12D^. Comparison of the docked poses of **11** (Fig 1C) and **13** (Fig 6B) suggests a potential rationale for the observed differences in binding affinity. The benzene ring of compound **11** is involved in a T-shaped π-stacking interaction with the side chain of Y71, which is replaced by the much smaller methyl in **13**. This suggests a critical role for the phenyl ring on the pyrimidine core for potency, providing a useful clue for future optimization efforts.

**Figure 5.**
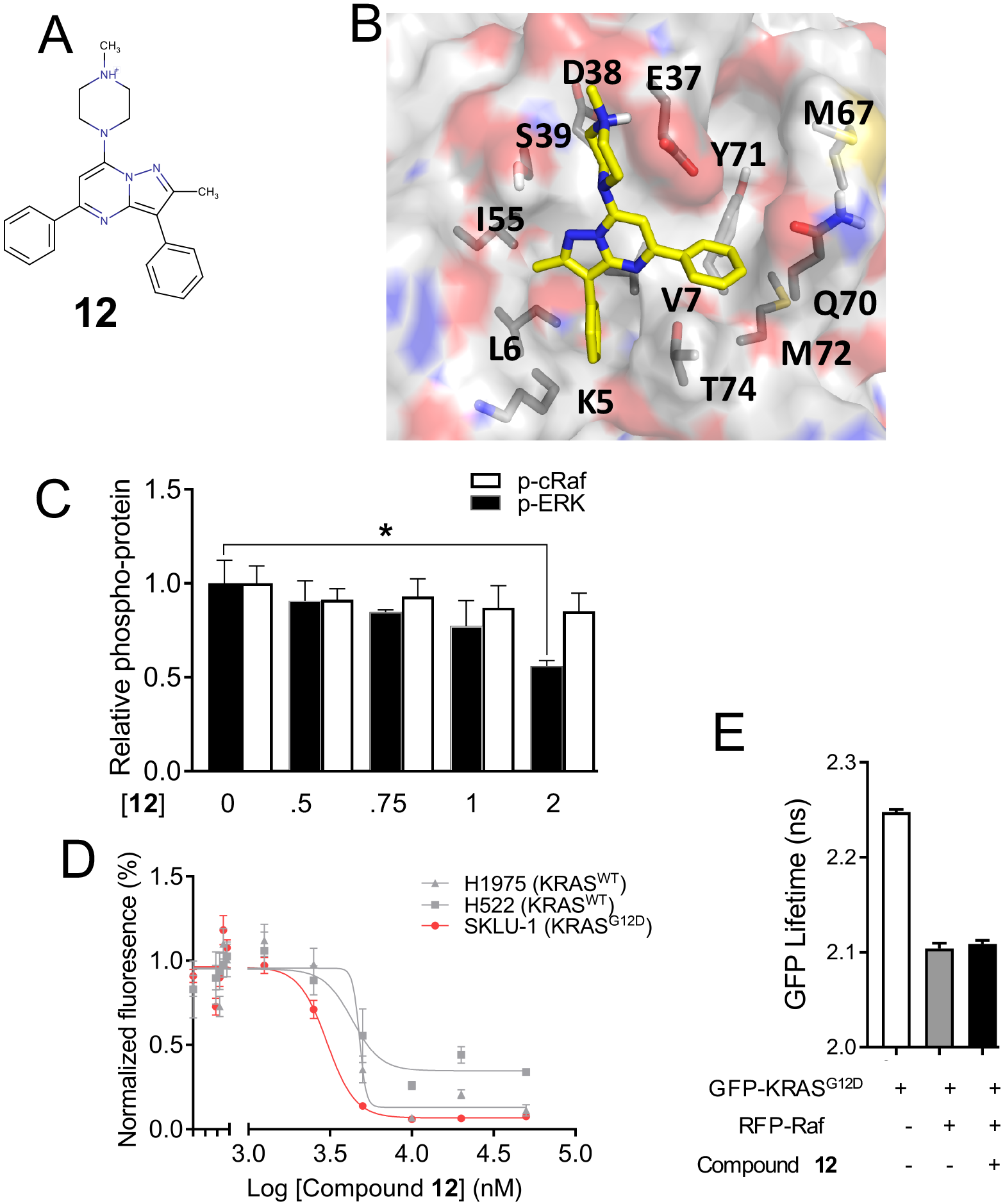
Potential role of the piperazineethanol moiety on compound 11 for abrogating effector binding. **(A**) Chemical structure of compound **12**, an analogue of **11** lacking the terminal hydroxymethyl functional group. **(B)** Predicted binding pose of compound **12**. (**C**) p-ERK and p-cRaf levels in BHK cells expressing KRAS^G12D^ treated with indicated concentrations of **12** or vehicle. **(D)** Proliferation profile of lung cancer cells upon treatment with increasing concentration of compound **12**, monitored by a CyQuant assay. Data are averages over three independent experiments and error bars represent standard error. **(E)** GFP fluorescence lifetime from FLIM-FRET using cells expressing GFP-KRAS^G12D^ alone or together with RFP-cRaf and with or without treatment with 2 μM compound **12**.

We then used fluorescence polarization and pull-down assays to test the functional implication of the modification in **13** relative to the parent compound **11**. Fig 6D shows that 20 μM of **13** disrupts the interaction of KRAS with GST-Raf^RBD^ as effectively as the parent compound. Our pull-down assay led to the same conclusion: **13** disrupts KRAS^G12D^-Raf^RBD^ interaction (Fig 6E). These results demonstrate that modifications can be made on the pyrazolopyrimidine core to optimize for potency without compromising effect on effector binding. This conclusion is supported by our structural analysis of the predicted ligand/KRAS complexes (Figs 1C and 6B), which shows that **11** and **13** likely make identical contacts with residues at the effector-binding region via their piperazine ring and especially the piperazineethanol group. This is important because, as we have shown using **12**, modification in this part of the ligand causes loss of effect on Raf binding. We then wondered if interaction with switch 2 residues or lack thereof may play a role in nucleotide release, because the conformation of many switch 2 residues, such as Y71 and Y64, differs between free and GEF-bound RAS (54, 55). To test this, we measured the intrinsic and SOS-dependent rates of labeled-GDP release in the absence and presence of **13**. We found that, indeed, replacing the benzene ring on the pyrimidine core by methyl dramatically altered the effect on nucleotide release. Whereas **11** had no effect on intrinsic and only modestly decreased the rate of SOS-mediated nucleotide release (Fig S5), **13** dramatically increased both rates (Fig 6F). This result suggests that interaction with switch 2 residues including Y71 may determine how a p1-bound ligand affects GEF activity. The results also provided strong support for the reliability of the predicted ligand-KRAS complex structures, and offered a viable route for additional modifications in future optimization efforts.

**Figure 6.**
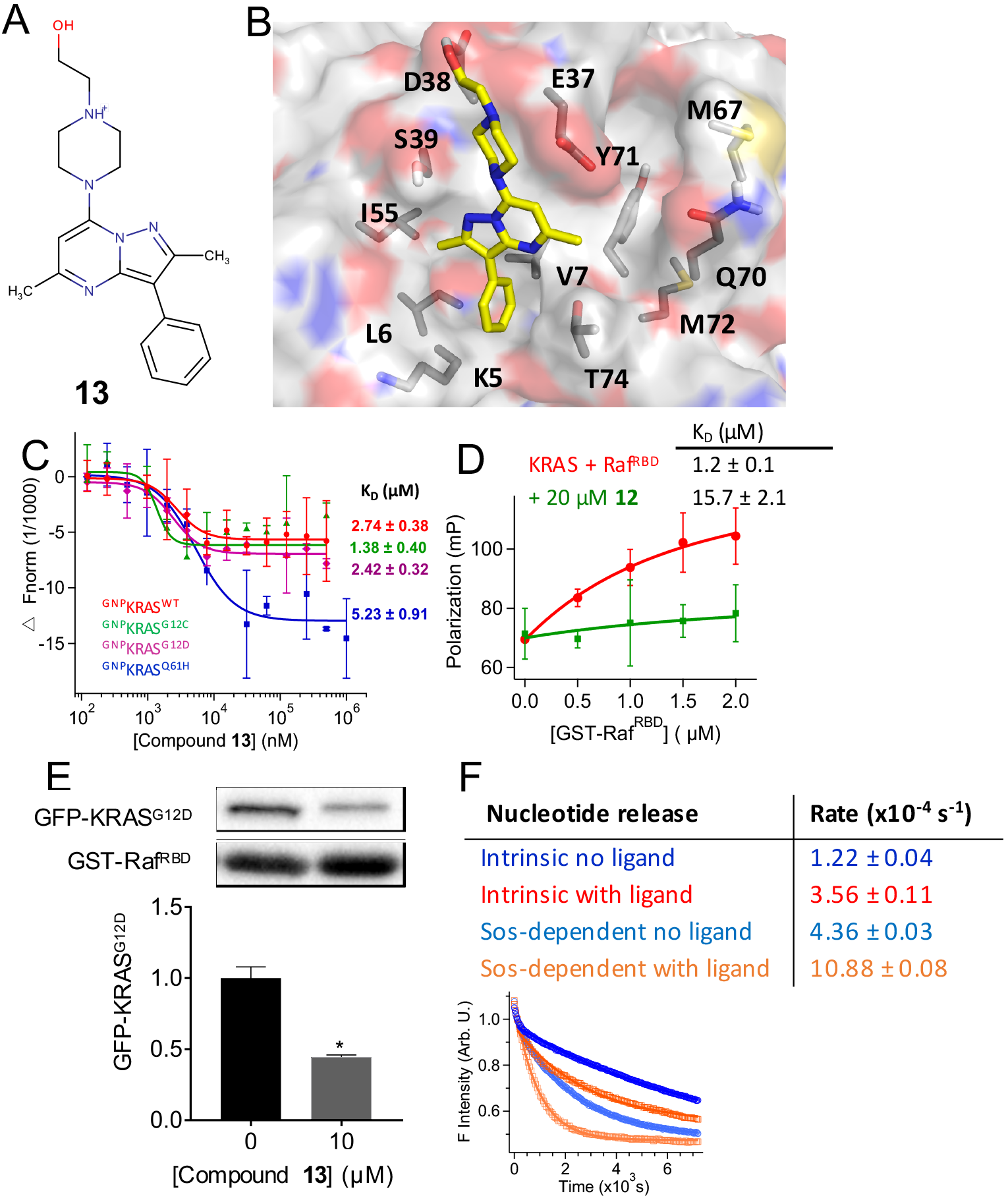
Interaction with switch 2 residues modulate exchange factor activity. **(A)** Chemical structure of compound **13**, an analogue of **11** without a benzene on the pyrimidine core. **(B)** Predicted binding pose of **13**. **(C)** Fluorescence intensity and K_D_ from MST experiments on KRAS mutants (see legend of Fig 2 for details). **(D)** Fluorescence polarization of ^BGTP-γ-S^KRAS (0.5 μM) with increasing concentration of GST-Raf^RBD^ in the absence (red) and presence (green) of 20 μM 13. **(E)** Amount of GFP-KRAS^G12D^ pulled down by GST-Raf^RBD^ after treatment of whole cell lysates with 10 μM of **13** (representative Western blots shown at the top). An equal volume of lysates was used and the data are normalized to GST-RBD and DMSO control. **(F)** Intrinsic (red) and SOS-mediated (blue) nucleotide release rates in a mixture of 0.5 μM KRAS (and SOS), 100 μM GTP and 0 or 50 μM of **13** (top) derived from changes in fluorescence intensity during the reaction KRAS^BGDP^ + GTP → KRAS^GTP^ + BGDP (bottom). Rates were calculated using single exponential fits starting a 120 s.

## Concluding Discussion

Finding a direct inhibitor of KRAS remains a major challenge in the search for cancer therapy. Previous attempts at preventing membrane binding of KRAS by farnesyl transferase inhibitors failed in clinical trials. More recent efforts focused on the dynamics of RAS revealed allosteric pockets suitable for binding of small molecules (32, 35). Several small-molecule ligands that bind to some of these pockets and disrupt interaction with GEFs or effectors have been discovered (21–25). However, thus far none of these ligands have led to a viable lead compound. In the current work, we combined MD simulation to generate a KRAS conformation with open pocket p1 and virtual screening to identify potential hits, followed by biophysical and cell biological experiments for validation. We have discovered a novel high-affinity KRAS inhibitor, compound **11**, that has unique structural features. Compound **11** (2-[4-(8-methyl-3,9-diphenyl-2,6,7-triazabicyclo[4.3.0]nona-2,4,7,9-tetraen-5-yl)piperazin-1-yl]ethanol) is drug-like (drug-likeness = 4.1) and somewhat polar with 6 hydrogen bond donors and 2 acceptors (cLogP = 0.87). It has a pyrazolopyrimidine core rather than an indole or imidazole ring typical in published ligands. Also, **11** is relatively large (415 Da) with its pyrazol ring methylated and benzylated and its pyrimidine ring beta-modified by benzene and 1-piperazineethanol. This allowed it to make more extensive predicted contacts with KRAS p1 residues than is common in most of the published ligands (Fig 1C). **11** selectively binds to ^GTP^KRAS with submicromolar affinity but not to GDP-bound KRAS, nor to GDP- and GTP-bound NRAS or HRAS (Fig 1, S3). It inhibits MAPK signaling (Fig 3) and proliferation of mutant KRAS-expressing but not HRAS-expressing cancer cells (Fig 4). Moreover, we used fluorescence polarization, pull-down and FLIM-FRET assays to demonstrate that compound **11** inhibits MAPK signaling primarily by abrogating interaction with effector proteins (Fig 2), in contrast to many published KRAS ligands that mainly affect GEF activity (22–24). At high concentration, **11** exhibits a small effect on intrinsic and GEF-catalyzed guanine nucleotide exchange rates (Fig S5), but the effect is too small at concentrations used in the cell-based assays to explain the significant inhibitory activity of the compound. For example, there is a maximum of ~5% reduction in the rates of both intrinsic and SOS-mediated nucleotide release or exchange reactions at 1 μM of **11**. In contrast, the p-ERK levels dropped by about 50% after a 3 h treatment using the same concentration of compound (Fig 3).

The above conclusions are also supported by data from comparative analyses of compound **11** and its analogues **12** and **13**. Compound **13** retains the effect of the parent compound on Raf binding even though it has a weaker (low μM) affinity for KRAS. Intriguingly, **13** accelerates both intrinsic and SOS-mediated rates of nucleotide release, in contrast to **11** which has no effect on the intrinsic and only modestly decreases the SOS-mediated reaction rate. Compound **12** has no effect on KRAS/Raf interaction and displays some inhibitory activities via an unknown mechanism. The distinct behavior of the derivatives and the parent compound, especially **11** and **13** for which we have data for direct KRAS binding, suggest altered protein-ligand interactions. We propose that the piperazineethanol group interacts with switch 1 of KRAS and plays a critical role in abrogating effector binding, whereas the potentially switch 2-interacting nonpolar moieties attached to the pyrazolopyrimidine core modulate GEF activity and contribute to high-affinity binding. These insights provide ideal starting points for further optimization of our highly promising lead compounds.

## Acknowledgment

We thank current and former members of the Gorfe and Hancock laboratories for illuminating discussions; Prof. X. Cheng for help with MST and for providing fluorescent-labeled Raplb; Prof. S. Cunha for help with pull-down assays. We gratefully acknowledge Prof. C. Mattos (Northeastern University) for purified GNP-bound KRAS samples; Prof. Sharon Campbell for HRAS and NRAS samples; Prof. J. Putkey and X. Wang (University of Texas Health Science Center) for purified GDP-bound KRAS samples and technical advices. We thank Dr. Jeffrey Myers of The University of Texas MD Anderson Cancer Center for providing the oral cancer cell lines. This work was supported by the Cancer Prevention and Research Institute of Texas (CPRIT grant No DP150093). K.J.C was supported in part by NIH (grant No R00CA188593).

## Author Contributions

A.A.G. designed and oversaw the project; M.J.M. performed high throughput virtual screening and cell signaling and proliferation assays; C.V.P. performed biophysical assays; P.P. performed MD simulations and structure analysis; M.J.M., K.J.C. and Y.Z. performed FLIM-FRET assays; L.H., A.K.G. and A.N. performed additional cell-based assays; K.J.C., D. vdH., J.F.H. and Y.Z contributed to the design and interpretation of cell assays; M.J.M., C.V.P., P.P., K.J.C. and A.A.G. wrote the paper. All authors contributed reagents/analysis tools, interpretation and critically reviewed the manuscript.

## Data availability statement

The data that support the findings of this study including MD-derived coordinates of KRAS used for docking are available from the corresponding author upon reasonable request.

## Conflict of interest statement

The authors declare no conflict of interest.

---

**Figure S1.**
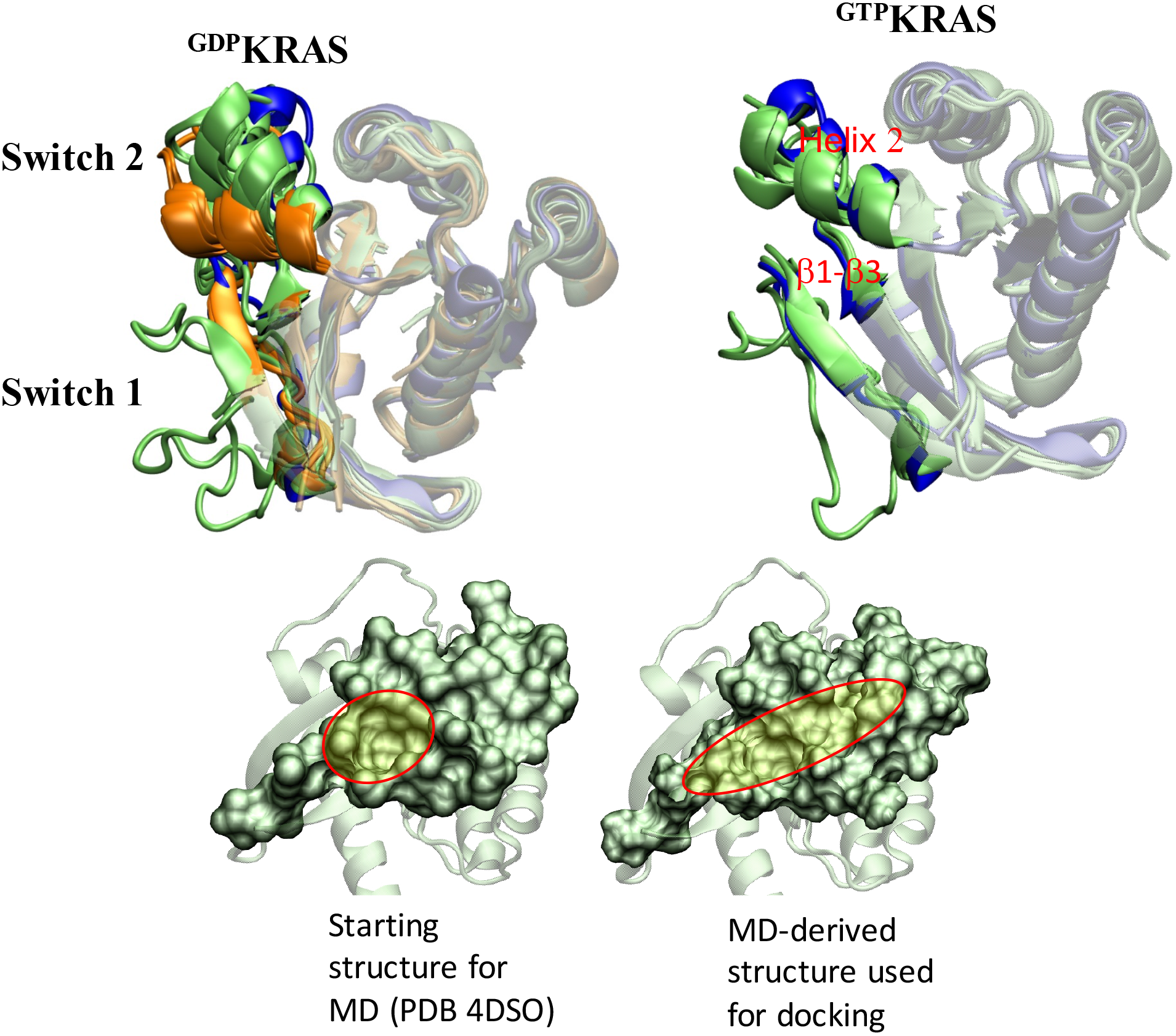
(Top) Multiseq structural alignment, using VMD (1), of the MD-derived conformer used for docking (blue) with 19 GDP-(5F2E, 4TQA, 4Q03, 40BE, 4M1T, 4LYJ, 4LV6, 4EPY, 4EPR, 6ASE, 6ARK, 5WHD, 5WHA, 5W22, 5VBM, 5V9U, 5V9L, 5V6V, 5UQW) and 4 GTP-or GTP analogue-bound KRAS X-Ray structures (5UK9, 5USJ, 4DSO, 6BP1). KRAS structures co-crystalized with proteins or peptides were not considered; all others have been included. Nearly half (eight) of the GDP structures contain covalent ligands targeting KRAS^G12C^ in which pocket p1 between switch 2 and the β1-β3 region is narrowed (orange). Based on the orientation of helix 2, three groups of conformations can be observed in the GDP structures, two of which also are sampled by the GTP structures. (Bottom) Displacement of helix 2 toward helix 3 and side chain reorganizations expanded pocket p1 (described by residues 5-7, 37-39, 50-56, 57-75 shown in surface representation in green; the expansion of a small hydrophobic pocket, originally occupied by benzamidine, during the MD simulation is highlighted in light green and circled in red).

**Figure S2.**
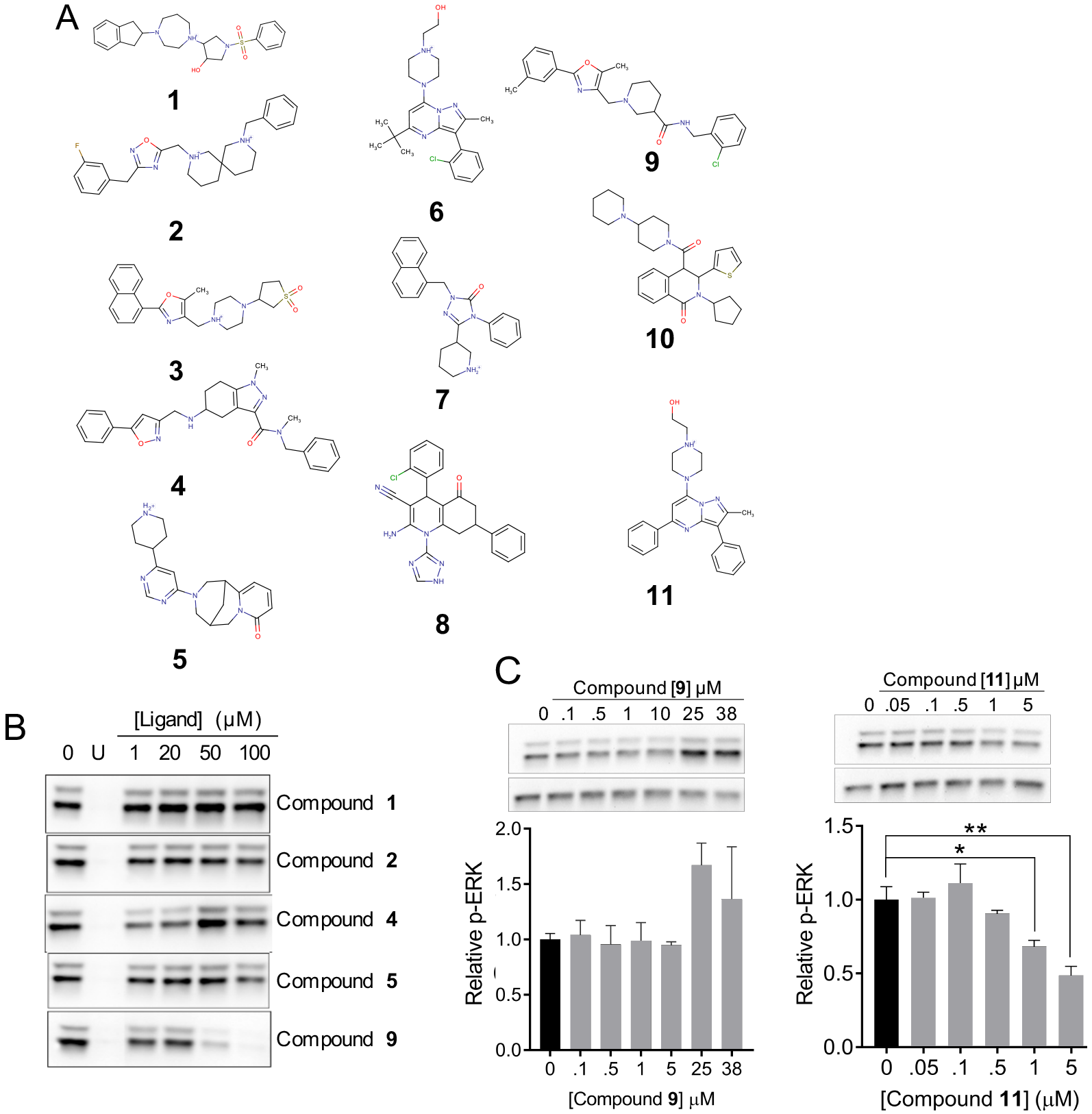
In silico prediction and initial experimental characterization of putative KRAS inhibitors. **(A)** Chemical structure of eleven computationally predicted hits (compounds **1** through **11**) selected for experimental testing in cell-based assays. **(B)** Western blots showing levels of phosphorylated ERK1/2 (p-ERK) in BHK cells ectopically expressing GFP-KRAS^G12D^ treated with vehicle (DMSO), a positive control U (the MEK inhibitor U0125) or compound at the indicated four concentrations. Five examples were shown illustrating cases where the ligand has no effect (compounds **1** and **2**), increases p-ERK levels (compound **4**) or decreases p-ERK levels (compounds **5** and **9**). This assay was used as a primary screen to quickly assess the potential of a ligand to qualitatively affect the MAPK pathway; for the sake of efficiency, we did not measure total protein. **(C)** Representative Western blots (top: p-ERK (upper) and total GFP-KRAS^G12D^ (lower), the latter serving as loading control), and their quantification (bottom) for compounds **9** and **11** for which the Western analysis was repeated in an expanded concentration range. The reduction in p-ERK levels of compound **11**-treated cells is not due to reduction in total ERK levels, as shown in Fig 3 of the main text. Data are averages over three independent experiments and error bars represent standard error. * = p < 0.02; ** = p < 0.005.

**Figure S3.**
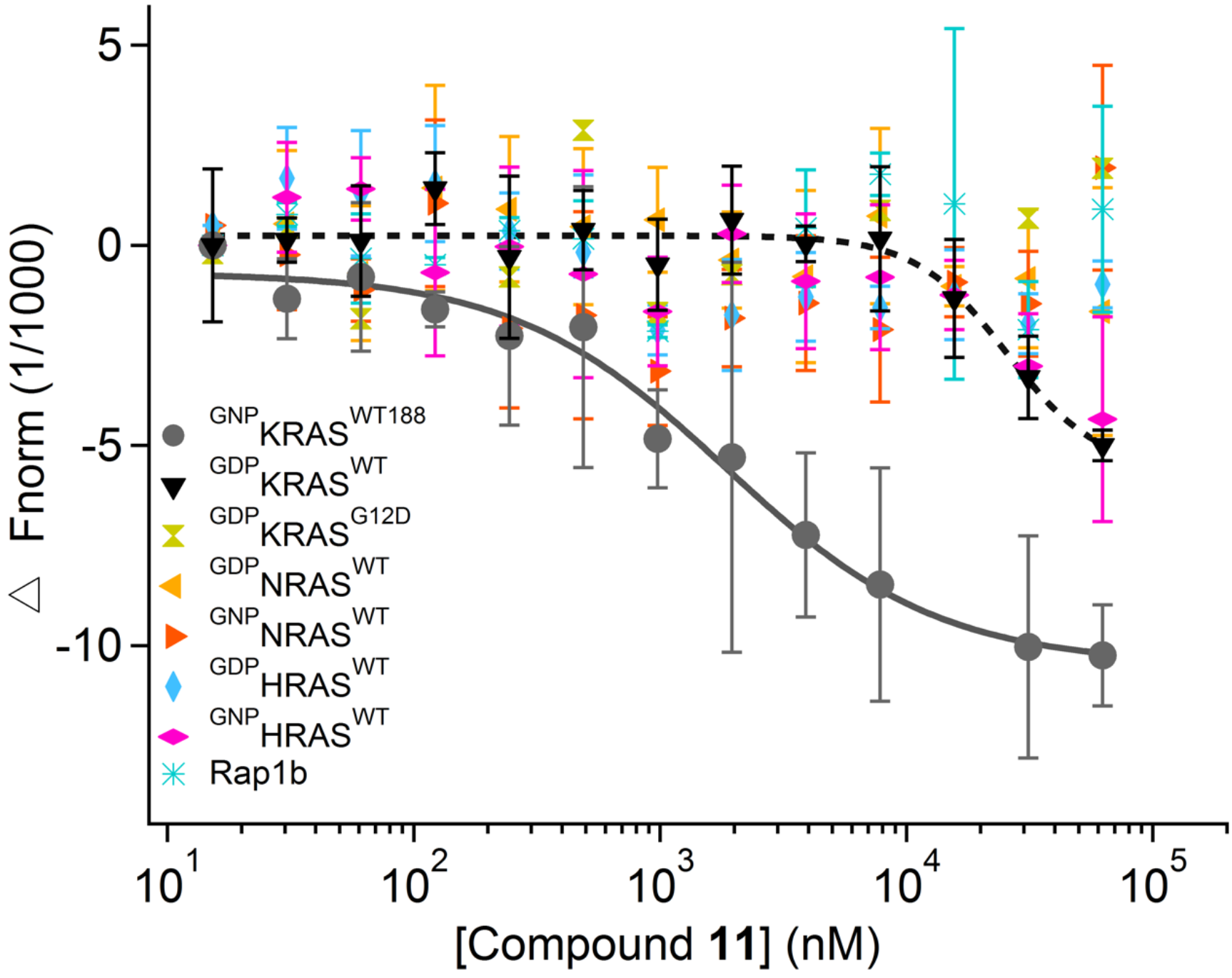
Microscale thermophoresis (MST) profiles monitoring the binding of compound **11** to the catalytic domain (residues 1-166) of ^GDP^KRAS^WT^ (black), ^GDP^KRAS^G12D^ (green), ^GDP^NRAS^WT^ (light orange), ^GNP^NRAS^WT^ (orange), ^GDP^HRAS^WT^ (light blue), ^GNP^HRAS^WT^ (pink), and Rap1b (cyan), as well as the full-length (residues 1-188; non-lipidated) ^GTP^KRAS^WT188^ (grey). MST profile of ^GDP^KRAS^WT^ (black) shows the onset of potential binding at > 20 μM compound **11** (^GNP^HRAS^WT^ exhibits a somewhat similar trend but the data is noisy). Note the weaker affinity for the full-length wild type protein relative to the isolated catalytic domain (estimated K_D_ of 1.9 ± 0. 4 versus ~0.3 (Fig 1 main text)). This suggests that the conformation of the full-length construct in solution is likely different from the isolated catalytic domain or cellular full-length KRAS (e.g., the p1 pocket could be partially occluded by the HVR in solution, which in the cell would have been engaged by the plasma membrane). Therefore, unless explicitly stated otherwise, we refer to the isolated catalytic domain throughout this manuscript when describing biophysical measurements.

**Figure S4.**
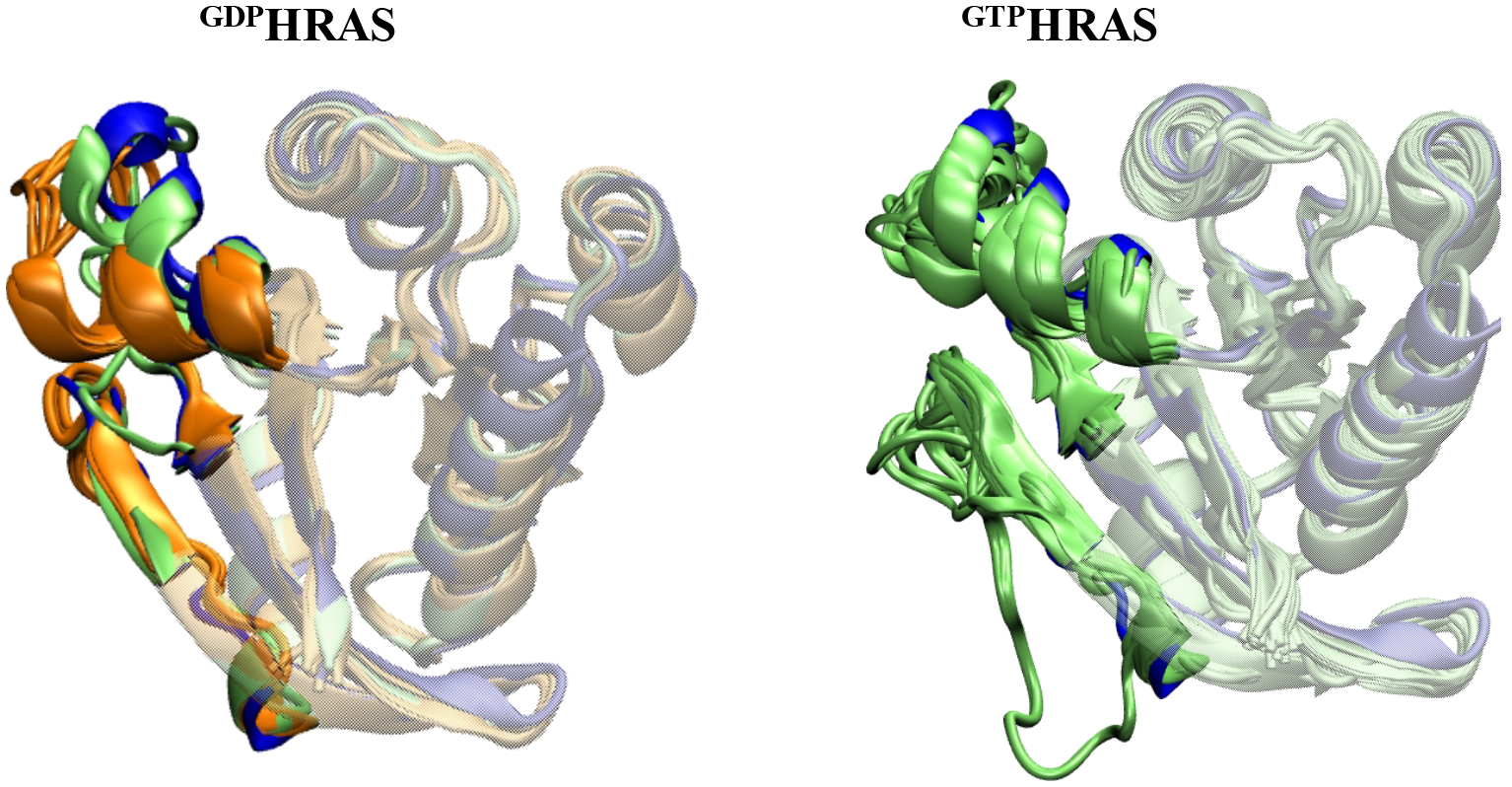
Alignment of the MD-derived KRAS conformer used for docking (blue) with 9 GDP-(1XJ0, 1ZVQ, 2CE2, 2CLD, 2QUZ, 3L05, 4L9S, 4Q21, 5VBE) and 27 GTP- or GTP-analogue-bound HRAS X-ray structures (1ZW6, 2CL6, 2CL7, 2EVW, 3I3S, 3K8Y, 3K9N,3KKM, 3KKN, 3RRY, 3TGP, 4DLR, 4DLS, 4DLT, 4EFN, 4XVQ, 5B2Z, 5B30, 5VNZ, 5WDP, 5WDQ, 5×9S, 6Q21). HRAS structures co-crystalized with proteins or peptides were not considered; all others have been included. As in KRAS, the orientation of helix 2 modulates the accessible space between the β1-β3 region and switch 2 but there is no obvious difference between the KRAS and HRAS structures at the global level. There aren’t as many experimental structures for NRAS.

## Effect of compound 11 on intrinsic and GEF-dependent nucleotide release and exchange reactions

Figure S8 shows time-dependent decreases and increases of fluorescence intensity as a labeled-nucleotide dissociates from and binds to KRAS^WT^, respectively. Compound **11** slightly reduced the rates of both intrinsic and SOS-mediated nucleotide exchange reactions, as well as the SOS-dependent (but not intrinsic) release of labeled-GDP. In particular, **11** decreased the intrinsic rate of nucleotide exchange by ~10-fold at >10 μM (Fig S5, top-left), but it has no effect on intrinsic nucleotide release (Fig S5, bottom-left). The latter is consistent with our observation from MST that **11** does not bind to ^GDP^KRAS^WT^ with high affinity. Since GTP hydrolysis is unlikely to occur within the timescale of our experiments (2, 3), a plausible interpretation of the former would be compromised GTP loading. This is possible if, for example, the ligand binds to the nucleotide free ‘transition state’ conformation of KRAS and induces reorganization of active site residues. This is supported by the fact that **11**’s effect on the rate of nucleotide exchange is significantly smaller (only a 1.1-fold decrease, Fig S5 top-right). SOS stabilizes nucleotide free RAS in an open active site conformation (4, 5), which allows for faster expulsion and rebinding of GTP or GDP.

**Figure S5.**
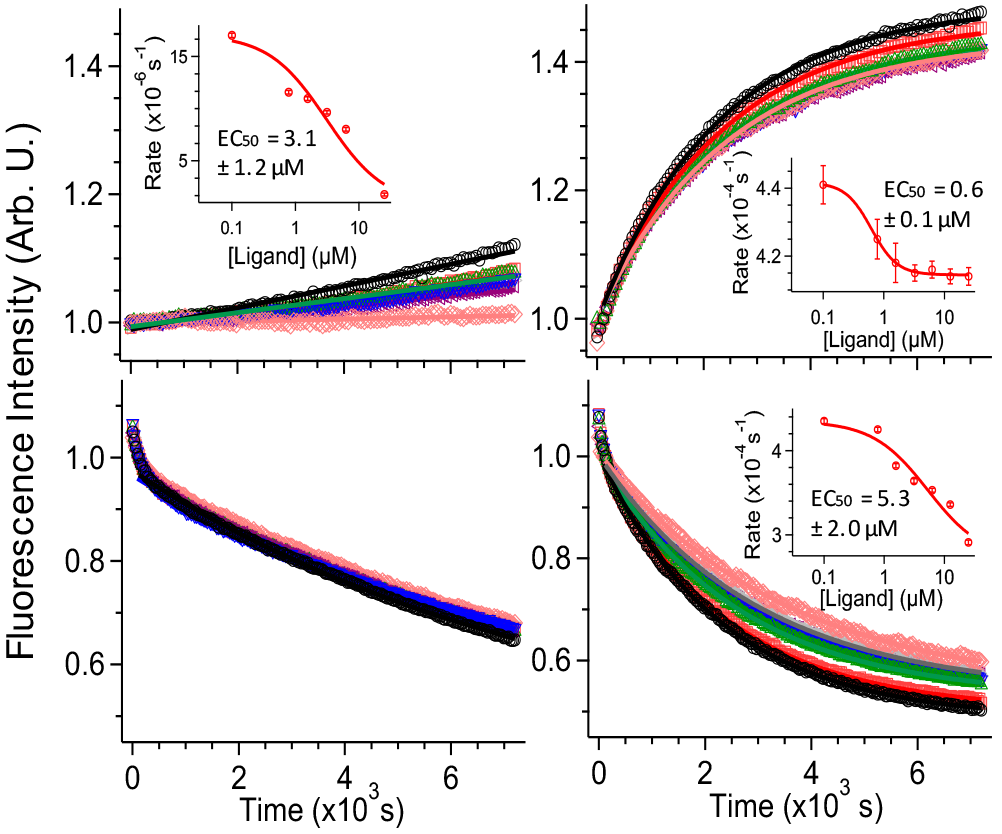
Effects of compound 11 on intrinsic and SOS-mediated nucleotide exchange and release. (**Top**) Intrinsic (left) and SOS-mediated (right) rates for the nucleotide exchange reaction KRAS^GDP^ + BGTP → KRAS^BGTP^ + GDP in a mixture of 0.5 μM each of KRAS, BGTP, and SOS. (**Bottom**) Intrinsic (left) and SOS-mediated (right) rates of the nucleotide release reaction KRAS^BGDP^ + GTP → KRAS^GTP^ + BGDP. Concentrations of KRAS and SOS were 0.5 μM and that of GTP was 100 μM. Ligand concentrations: 0 (circle), 0.78 (square), 1.56 (triangle), 3.12 (inverted triangle), 6.25 (left-sided triangle), 12.5 (right-sided triangle) and 25 μM (diamond). Intensities were normalized with respect to the value at 120 s. Linear or single exponential fits, starting from 120 s, were superimposed as solid lines. Inset: Rates as a function of ligand concentration.

Plotting the measured rates as a function of the total ligand concentration yielded additional insights into the effect of **11** in the intrinsic and GEF-catalyzed enzymatic activity of KRAS. We obtained an estimated EC_50_ of 3.1 ± 1.2 μM from the intrinsic nucleotide exchange assay, and a similar value of 5.3 ± 2.0 μM from the SOS-mediated GDP release assay. These values are about 10-times larger than the K_D_ of **11** for ^GTP^KRAS^WT^, suggesting a potentially weaker binding to the nucleotide free state assuming, as argued above, that the observed effects on enzymatic activity are at least in part a result of binding to nucleotide free KRAS. Intriguingly, we obtained EC_50_ = 0.6 ± 0.1 μM from the SOS-dependent nucleotide exchange measurements, a value very close to the K_D_ of **11** for ^GTP^KRAS^WT^. This may reflect binding to the GTP-bound KRAS at the allosteric site of SOS. We note that all of the effects we observed on reaction rates are much smaller than those in KRAS-Raf interaction. Nonetheless, they are statistically significant and dose-dependent, suggesting that compound **11** may modulate KRAS activation through multiple mechanisms, with the dominant effect being on effector binding.

